# Genome Sequences of *Populus tremula* Chloroplast and Mitochondrion: Implications for Holistic Poplar Breeding

**DOI:** 10.1101/035899

**Authors:** Birgit Kersten, Patricia Faivre Rampant, Malte Mader, Marie-Christine Le Paslier, Rémi Bounon, Aurélie Berard, Cristina Vettori, Hilke Schroeder, Jean-Charles Leplé, Matthias Fladung

## Abstract

Complete *Populus* genome sequences are available for the nucleus (*P. trichocarpa;* section *Tacamahaca*) and for chloroplasts (seven species), but not for mitochondria. Here, we provide the complete genome sequences of the chloroplast and the mitochondrion for the clones *P. tremula* W52 and *P. tremula* x *P. alba* 717–1B4 (section *Populus*). The organization of the chloroplast genomes of both *Populus* clones is described. A phylogenetic tree constructed from all available complete chloroplast DNA sequences of *Populus* was not congruent with the assignment of the related species to different *Populus* sections. In total, 3,024 variable nucleotide positions were identified among all compared *Populus* chloroplast DNA sequences. The 5-prime part of the LSC from *trnH* to *atpA* showed the highest frequency of variations. The variable positions included 163 positions with SNPs allowing for differentiating the two clones with *P. tremula* chloroplast genomes (W52 717–1B4) from the other seven *Populus* individuals. These potential *P. tremula*-specific SNPs were displayed as a whole-plastome barcode on the *P. tremula* W52 chloroplast DNA sequence. Three of these SNPs and one InDel in the *trnH-psbA* linker were successfully validated by Sanger sequencing in an extended set of *Populus* individuals. The complete mitochondrial genome sequence of *P. tremula* is the first in the family of *Salicaceae*. The mitochondrial genomes of the two clones are 783,442 bp (W52) and 783,513 bp (717–1B4) in size, structurally very similar and organized as single circles. DNA sequence regions with high similarity to the W52 chloroplast sequence account for about 2% of the W52 mitochondrial genome. The mean SNP frequency was found to be nearly six fold higher in the chloroplast than in the mitochondrial genome when comparing 717–1B4 with W52. The availability of the genomic information of all three DNA-containing cell organelles will allow a holistic approach in poplar molecular breeding in the future.

## Introduction

Chloroplasts and mitochondria are organelles within living cells, with the former occurring only in green land plants and in algae. Chloroplasts are very important for the conversion of sunlight into chemical energy, while the main function of mitochondria is to convert the energy from food into chemical energy. Plastids and mitochondria arose through endosymbiotic acquisition of formerly free-living bacteria, explaining the presence of own genomes [1–3]. During more than a billion years of subsequent concerted evolution, the three genomes of plant cells (nuclear, plastid and mitochondrial) have undergone dramatic structural changes to optimize the expression of the compartmentalized genetic material [1]. Prokaryotic genes disappeared from the organellar genomes and were integrated into the nuclear genome or were lost in a lineage-specific manner [1,4,5]. Nuclear-cytoplasmatic co-evolution can cause genetic incompatibilities that contribute to the establishment of hybridization barriers, ultimately leading to the formation of new species [1].

In most seed plants, both chloroplasts and mitochondria are inherited maternally [6], while nuclear genetic information is inherited bi-parentally [7]. There is only one allele per cell and per organism in the genetic information of the organelle DNA. The double-stranded organelle DNA is present in many copies per cell, leading to the convenient situation that organelle DNA can be retrieved relatively easily from low-quantity and/or degraded DNA samples. These advantages are reflected in the recommendation of the Barcode of Life Consortia to apply molecular markers, mainly based on organelle DNA, for the genetic differentiation of all eukaryotic species [8]. For genetic differentiation of most animals, the mitochondrial encoded cytochrome oxidase subunit I (*COX1*) gene sequence is the default barcode region [9]. For genetic differentiation of vascular plants, molecular markers are used mainly based on DNA variations in two chloroplast regions (*rbcL* and *matK*) as a two-locus barcode [10,11]. In addition, chloroplast DNA (cpDNA) and mitochondrial DNA (mtDNA) molecular markers can be applied in inter-specific hybrids to determine the maternal cross partner and to unravel the direction of the cross [12–14]. Chloroplast markers are also largely used to study the phylogeography of many plant species including trees (*e.g*., [15,16]).

For the development of cp- or mtDNA molecular markers, the availability and cross-species comparability of complete organelle genome sequences is particularly beneficial. In the last ten years, the number of complete plant organelle genome sequences has increased explosively along with the development of the next generation sequencing (NGS) tools. According to the Organelle Resources at the NCBI database ([17]; last page view November 9, 2015), 622 records are listed for complete chloroplast genome sequences and 303 records for mitochondrial genome sequences of land plants.

The size of most plant chloroplast genomes ranges from 150 kb to 160 kb, with few less than 100 kb, mostly organized as circular molecule with highly conserved structure. The cpDNA is often divided into four parts, including a large single copy (LSC) and a small single copy (SSC) region, interrupted by a pair of large, about 25 kb long, inverted repeats (IRa and IRb). The plastome in photosynthetic plants comprises 70 (gymnosperms) to 88 (liverworts) protein coding genes and 33 (most eudicots) to 35 (liverworts) structural RNA genes, totaling 100–120 unique genes [18].

In contrast, the mitochondrial genomes of plants vary greatly in size (mostly ranging from 200 to 600 kb), gene content and gene order [19]. The smallest known mitochondrial genome of land plants is of a size of approximately 65.9 kb (*Viscum scurruloideum*) and the largest of about 11.3 Mb (*Silene conica;* [17]; last page view November 9, 2015). Gene content also varies considerably among species, ranging from 32 to 67 genes, some of which are encoded in discontinuous fragments that require trans-splicing [19].

The genus *Populus* consists of about 29 different species, classified in six different sections (*Populus* (formerly *Leuce*)*, Tacamahaca, Aigeiros, Abaso, Turanga* and *Leucoides* [20]). Most *Populus* species are dioecious, although reports of hermaphroditism have been published (reviewed in [21]). Due to its small genome size, huge genomic resources and easy biotechnological handling ability, this genus has become a model species for tree genomics [22], resulting in the publication of the full genome of western black cottonwood, *P. trichocarpa* [23]. Only few of the *Populus* species are of high scientific and economic importance, and serve as source material for breeding purposes [24,25]. Important characteristics of some inter-specific hybrids are high growth rates and a broad applicability ranging from wood and paper to energy production [26]. Nowadays, breeding of *Populus* is intensified worldwide due to the application of biotechnological methods and the availability of huge genomic resources [27,28]. Superior clones of various *Populus* species have been developed and are commonly used for biomass production in short rotation plantations. Some poplar clones are easily accessible for genetic transformation, and many transgenic poplar lines have been transferred to the field and tested for commercial application [29].

For the genus *Populus*, complete cpDNA genome sequences have been made available for *P. trichocarpa* (GenBank accession number EF489041; [23]), *P. alba* (AP008956; [30]), *P. balsamifera* (KJ664927; [31]), *P. cathayana* (KP729175), *P. euphratica* (KJ624919; [32]), *P. fremontii* (KJ664926; [31]), and *P. yunnanensis* (KP729176). The related genome sizes range from 155,449 bp (*P. cathayana*) to 157,446 bp (*P. fremontii*) and the number of annotated proteins from 76 (*P. cathayana*) to 98 (*P. trichocarpa*).

For genetic differentiation of *Populus* species, several other species-specific cpDNA molecular markers have been developed in addition to the two official barcode markers [13,33,34]. No complete genome sequence for mitochondria has been published for any *Populus* species yet. Although Tuskan et al. [23] reported the assembly of a circular putative master molecule of 803 kb for the mitochondrial genome of *P. trichocarpa*, the related nucleotide sequence is not available so far. There is only a partial *P. trichocarpa* mitochondrial sequence of 230 kb available at GenBank (accession number KM091932; Boehm and Cronk, unpublished). This sequence has been declared as “unverified” by GenBank.

Only few attempts have been made to develop molecular markers inside the mitochondrial genome for species or population differentiation. Most of these approaches were based on the use of restriction fragment length polymorphisms in poplars [35,36]. Quite recently, also several mitochondrial SNP markers were made available allowing the genetic differentiation of some *Populus* species [12].

In this study, we provide the complete DNA sequence of the chloroplast genome of two clones from the section *Populus, P. tremula* W52 and *P. tremula* x *P. alba* 717–1B4, and compare the DNA sequences with those of all other chloroplast genomes available from the genus *Populus*. These analyses reveal comparative structural and functional information, broadening the knowledge base of *Populus* cpDNA and stimulating future diagnostic marker development.

Further and more importantly, we report the mitochondrial genomes for the two clones as the first complete mitochondrial DNA genome sequences of any *Populus* species, and even the first in the family of *Salicaceae*. The door is now open to apply genomic sequences of all three DNA-containing cellular compartments, thus stimulating molecular *Populus* breeding efforts. This will allow, for the first time, a holistic approach in a tree breeding program considering all genomic DNA-containing subcellular compartments.

## Materials and Methods

### Plant material

The male *P. tremula* clone W52 originates from a seed collection harvested from aspen trees in Gwardeisk (Russia; former Tapiau) in the 1920s/30s, and since then germinated seedlings were grown in Wedesbüttel, Meine, Germany. Since the early 1950s, shoots of W52 were grafted on unrelated *P. tremula* root stocks and cultivated in the clonal archive of the Thuenen Institute of Forest Genetics, Grosshansdorf, Germany [37].

INRA clone 717–1B4 is a female clone, originating from a cross between a female *Populus tremula* tree (tree # 5903) harvested on March 3, 1959 in the Parroy forest (Meurthe Moselle, France) and a male *Populus alba* tree (tree # 6072) harvested on February 22, 1960 close to the Pont du Gard (France). This hybrid was part of a breeding program initiated by Dr. Michel Lemoine in the 1960's [38]. Trees 5903 and 6072, installed in the nurseries of Nancy, were lost during storms in 1999 and vegetative copies of the parents are no longer available.

By employing established chloroplast PCR-RFLP markers to differentiate *Populus* species (primer sequences are given in S1 appendix), *i.e*. the *P. tremula*-specific marker *rpoC2-rpoC1* (*Mls*I) and the *P. alba*-specific marker *trnH-psbA* (Alw26I) [13,14], the maternal part in the 717–1B4 clone could be without doubt classified as *P. tremula* (S2 appendix; 717–1B4 provided a positive result with the *P. tremula-specific* marker and a negative result with the *P. alba*-specific marker). Our results were reinforced by recently developed mitochondrial molecular markers [12].

### Whole genome shot gun sequencing of W52 and 717–1B4

The Ion Torrent sequencing platform was used for the shotgun sequencing of total genomic DNA of the W52 sample using the Personal Genome Machine^®^ (PGM™) Sequencer (Life Technologies, USA). Total genomic DNA (100 ng) was sheared using the Ion Shear™ Plus Reagents and used for preparing the sequencing library according to the Ion Xpress^TM^ Plus gDNA Fragment Library kit (catalog number 4471252) following Ion Torrent PGM™ protocol (Life Technologies, USA). The resulting individual DNA library was quality checked and quantified using the Qubit^®^ 2.0 Fluorometer and the Qubit^®^ dsDNA HS Assay Kit following the manufacturer's specification (Life Technologies, USA). Following template amplification and enrichment on the Ion OneTouch™ 2 System (Ion OneTouch™ 2 Instrument for amplification and Ion OneTouch™ ES enrichment system, Life Technologies, USA) using the Ion PGM™ Template OT2 400 Kit (catalog number 4479878), the sample was loaded onto one PGM Ion 318™ Chip v2 and sequenced using the Ion PGM™ Hi-Q™ Sequencing Kit (catalog number A25592, Life Technologies, USA) according to manufacturer's protocol.

Ramets of INRA 717–1B4 were grown in the greenhouse. The youngest leaves were harvested, shock-frozen in liquid nitrogen and stored at −80°C. Frozen leaves were then ground with a pestle and mortar to a fine powder in liquid nitrogen. Total genomic DNA was extracted using the Dolezel-MATAB (mixed alkyltrimethylammonium bromide) method [39]. Genomic DNA was then prepared for the Illumina-MiSeq and Ion-Proton sequencing systems. The whole genome sequencing (WGS) DNA sequencing library for Illumina-MiSeq sequencing was created using the TruSeq DNA PCR-Free LT kit (Illumina). Briefly, sample preparation was performed with 2 μg of DNA using the low sample protocol. The mean fragment size was 550 bp. All enzymatic steps and cleaning steps, including fragmentation using AFA (Adaptive Focused Acoustics™) technology on focused-ultrasonicator E210 (Covaris), were performed according to the manufacturer's instructions. On board clusters generation and pair-end sequencing (2 × 250 bp sequencing by synthesis cycles) were performed on a MiSeq (Illumina) according to the manufacturer's instructions.

A WGS DNA sequencing library of 717–1B4 was prepared with the Ion Xpress™ Plus Fragment Library Kit for Ion-Proton sequencing and sequencing was performed as described above for W52.

### Bioinformatic analyses

If not stated otherwise, all bioinformatic analyses steps were performed using CLC Genomics Workbench (CLC GWB; v7.0.4; CLC bio, a QIAGEN Company, Aarhus, Denmark).

#### Assembly and annotation of the complete cpDNA sequences of W52 and 717-1B4

The chloroplast genome of 717–1B4 was generated first, as paired-end data were available only for 717–1B4. Sequencing reads from total genomic DNA of 717–1B4 (miSeq and IonTorrent) were trimmed and *de novo* assembled (methods described in detail in S3 appendix). Putative chloroplast contigs (larger than 1000 bp) were selected from the miSeq and IonTorrent contigs based on mean coverage and further refined by mapping trimmed miSeq and IonTorrent reads and subsequent consensus calling. During consensus calling, regions of low coverage were removed and contigs were split when including regions of low coverage. Conflicts were resolved by voting using quality scores.

Consensus chloroplast contigs larger than 300 bp were subjected to scaffolding using SeqMan Pro (v10.1.2.; DNA Star, Madison, USA; Pro assembly with default parameters, but increased match size of 40). Only the largest scaffold (128,565 bp) was kept, as this scaffold already included the LSC, a *collapsed* consensus of *IRb/IRa* and the SSC. The missing complete IRa was created from the *collapsed* consensus of *IRb/IRa* by calculating the reverse complement of this sequence. Using small stretches of IRa/IRb at both ends of the scaffold, the IRa sequence was then combined with the scaffold to get a complete cpDNA sequence. The 5-prime-end of the sequence was adjusted to the beginning of the LSC (starting with the reverse complement of the *trnH* gene).

Using the complete cpDNA sequence of 717–1B4 as a reference, the related W52 sequence was assembled by mapping the trimmed W52 reads (trimming of IonTorrent reads described in S3 appendix) to the reference and subsequent calling of a consensus sequence. During consensus calling, regions of low coverage were filled by Ns and conflicts were resolved by voting using quality scores. Gaps in the consensus were closed manually either by adding missing regions from W52 contigs (generated by an independent *de novo* assembly of IonTorrent reads according to S3 appendix) or by extending gap borders after mapping of W52 reads until gap closure. Trimmed W52 reads were mapped to the edited consensus sequence and the mapping was checked for breaks. These were treated manually by splitting the sequence at the breakpoint, extending both ends by mapped reads until a combination of overlapping ends was possible. New combined sequence regions were validated in a subsequent read mapping.

In the first run, the complete chloroplast sequences of W52 and 717–1B4 were automatically annotated using the CpGAVAS web server [40,41]. Parameters were set to default with the exception of the “minimal numbers of member genes/protein for a homologous group to be included as reference gene/protein sets for annotation” that was set to 10. Genes related to protein sequences that missed start codons or stop codons or included frameshifts based on the CpGAVAS prediction were re-annotated using DOGMA [42] or based on annotation of the related gene in the *P. alba* cpDNA sequence (GenBank accession number AP008956). Additional tRNA genes that include introns were predicted based on annotation of the related gene in the *P. alba* cpDNA sequence.

#### Assembly and annotation of the complete mtDNA sequences of W52 and 717-1B4

The trimmed miSeq and IonTorrent reads of 717–1B4 were cleaned from chloroplast reads by mapping to the 717–1B4 chloroplast sequence (0.99 identity; 0.99 overlap). Unmapped reads representing nearly chloroplast-free reads were extracted and *de novo*-assembled. Putative mitochondrial contigs larger than 1,000 bp were selected from the resulting contigs based on mean coverage and – in parallel – based on BLAST of all contigs against selected mitochondrial genome sequences available at NCBI [43]. After refinement of the contigs (as described for chloroplast contigs; see above), consensus contigs larger than 300 bp were subjected to scaffolding using SeqMan Pro (v10.1.2; Pro assembly with default parameters, but increased match size of 40). All scaffolds of sizes above 2,000 bp were kept and further extended by repeated mappings of the trimmed miSeq and IonTorrent reads followed by scaffold extensions. The complete mitochondrial sequence was manually finished by combining overlapping scaffold ends. After completing the mitochondrial genome of 717–1B4, the trimmed reads were mapped to identify regions of low coverage as well as regions of high coverage (due to additional unspecific mapping of very similar chloroplast-derived reads). Sequences of both types of regions were verified by Sanger sequencing of PCR amplicons of mtDNA of 717–1B4 as described below.

The mitochondrial genome sequence of W52 was generated by reference-based assembly based on the complete mtDNA sequence of 717–1B4 (as described above for the chloroplast sequence of W52).

Protein-coding genes, tRNA and rRNA genes were predicted in the complete mitochondrial genome sequence of W52, using the MITOFY webserver [44,45]. Intron-less genes were refined – if necessary – according to full ORFs predicted by ORF finder at NCBI [46]. Gene models of intron-including genes, especially trans-spliced genes that could not sufficiently be predicted using MITOFY were derived based on related protein sequences of other *Populus* species (available only for a few genes) or based on gene models in the related mitochondrial genome of *Ricinus communis* (GenBank accession number HQ874649). Additional putative protein-coding genes were predicted using ORF finder at NCBI [46]. The nucleotide sequences of all identified ORFs larger than 400 bp were analysed by BlastN versus *P. tremula* transcriptome assemblies at PopGenie v3 (page view November 4, 2015) [76,77]. Only ORFs with supporting transcript data (BlastN hit with at least 99.5% identity) were selected as putative protein-coding genes. The related genes were annotated as “unknown” and the related proteins as “hypothetical proteins”, because no putative function could be assigned to these proteins so far (BlastX versus NCBI nonredundant protein sequences).

BlastN analysis (CLC GWB) of the W52 mtDNA sequence versus the W52 cpDNA sequence (e-value: 1E^-18^; word size 25) was performed to identify genomic regions in the W52-mtDNA which show high similarity to genomic regions in the W52-cpDNA. Regions of the mitochondrial genome showing hits with at least >90% identity and at least 100 bp hit length were selected as chloroplast-like DNA regions in the mitochondrial genome of W52.

The mitochondrial genome of 717–1B4 was annotated by transferring the annotations from the W52 mitochondrial sequence to 717–1B4.

#### Identification of sequence variations between cp/mtDNA of 717–1B4 and W52

In order to identify DNA sequence variations in the chloroplast and mitochondrial genomes of 717–1B4 compared to W52, trimmed W52 reads were mapped to the complete chloroplast/mitochondrial genome sequence of 717–1B4 using default parameters but increased length fraction (95%) and similarity fraction (99%). SNPs and small InDels were identified using quality based variant detection (default parameters with minimum variant frequency of 90%).

#### Alignment of complete *Populus* cpDNA sequences, construction of a phylogenetic tree and SNP detection in the alignment

Complete cpDNA sequences of *P. tremula* W52 (GenBank accession number KP861984), *P. tremula x alba* 717–1B4 (KT780870), *P. trichocarpa* (EF489041), *P. alba* (AP008956), *P. balsamifera* (KJ664927), *P. cathayana* (KP729175), *P. euphratica* (KJ624919), *P. fremontii* (KJ664926), and *P. yunnanensis* (KP729176) were aligned together with the complete chloroplast sequence of *Salix interior* (NC_024681). Pairwise comparisons were run using the alignment file as an input. The best-fitting model for this alignment was selected prior to phylogenetic tree construction. A phylogenetic tree was constructed using UPGMA (unweighted pair group method with arithmetic mean) as construction method, general time reversible as nucleotide substitution model, including rate variations (number of substitution rate categories = 4; gamma distribution parameter = 1). A bootstrap analysis was performed with 100 replicates.

In order to identify SNPs and small InDels between all *Populus* chloroplast sequences, the alignment described in the previous section was repeated but without including the chloroplast sequence of S. *interior*. The alignment-FASTA was exported from CLC GWB and used as an input for the web tool SNiPlay (pipeline v2) [47,48] to run SNP/INDEL discovery (default parameters).

### PCR amplification and Sanger sequencing of specific cpDNA/mtDNA regions in *Populus*

Nucleotide sequences of the primers used for PCR are given in S1 appendix. PCR reactions were performed in 1×reaction buffer BD (provided together with Taq-polymerase by DNA Cloning Service, Hamburg, Germany), 1.8 or 2.0 mM MgCl_2_, 200 μM dNTP-Mix, 0.3 μM of each primer, 5% dimethyl sulfoxide, 1 U DNA polymerase and 50 to 100 ng DNA (in a total volume of 25 μL). The PCR program was started with an initial denaturation for 3 min at 94°C. Thirty-five to 40 PCR cycles followed, with 30–60 s at 94°C, 45 s at the respective annealing temperature, and 60 s-120 s (depending on the PCR product size) at 72°C. The reaction was completed by a final elongation step for 10 min at 72°C. Annealing temperatures were calculated on the basis of the primer sequences. Five μL of each PCR product were visualized on 1.2% agarose gel (120 V, 1:20 h) stained with the DNA fluorescence additive Roti-Safe Gel Stain (Carl Roth, Karlsruhe, Germany). The PCR products were purified with lithium chloride (5 μL LiCl), precipitated by adding 130 μL of absolute ethanol (overnight at −70°C), and Sanger-sequenced by StarSeq (Mainz, Germany).

## Results

### Organization of the *P. tremula* chloroplast genome and sequence variations between W52 and 717–1B4

The complete nucleotide sequence of the chloroplast genome of *P. tremula* was identified for the male *P. tremula* clone W52 (GenBank accession number KP861984) and the female *P. tremula* x *P. alba* clone INRA 717–1B4 (KT780870) based on whole genome shot gun sequencing of total genomic DNA by NGS.

Fig. 1 (middle circle) shows the chloroplast genome organization of *P. tremula* based on the cpDNA sequence of W52 (156,067 bp). The gene map was created by OrganellarGenomeDraw [49]. The genome includes a LSC of 84,367 bp and a SSC of 16,670 bp that are separated by a pair of inverted repeats, each of a length of 27,509 bp (IRa and IRb).

**Fig. 1.**
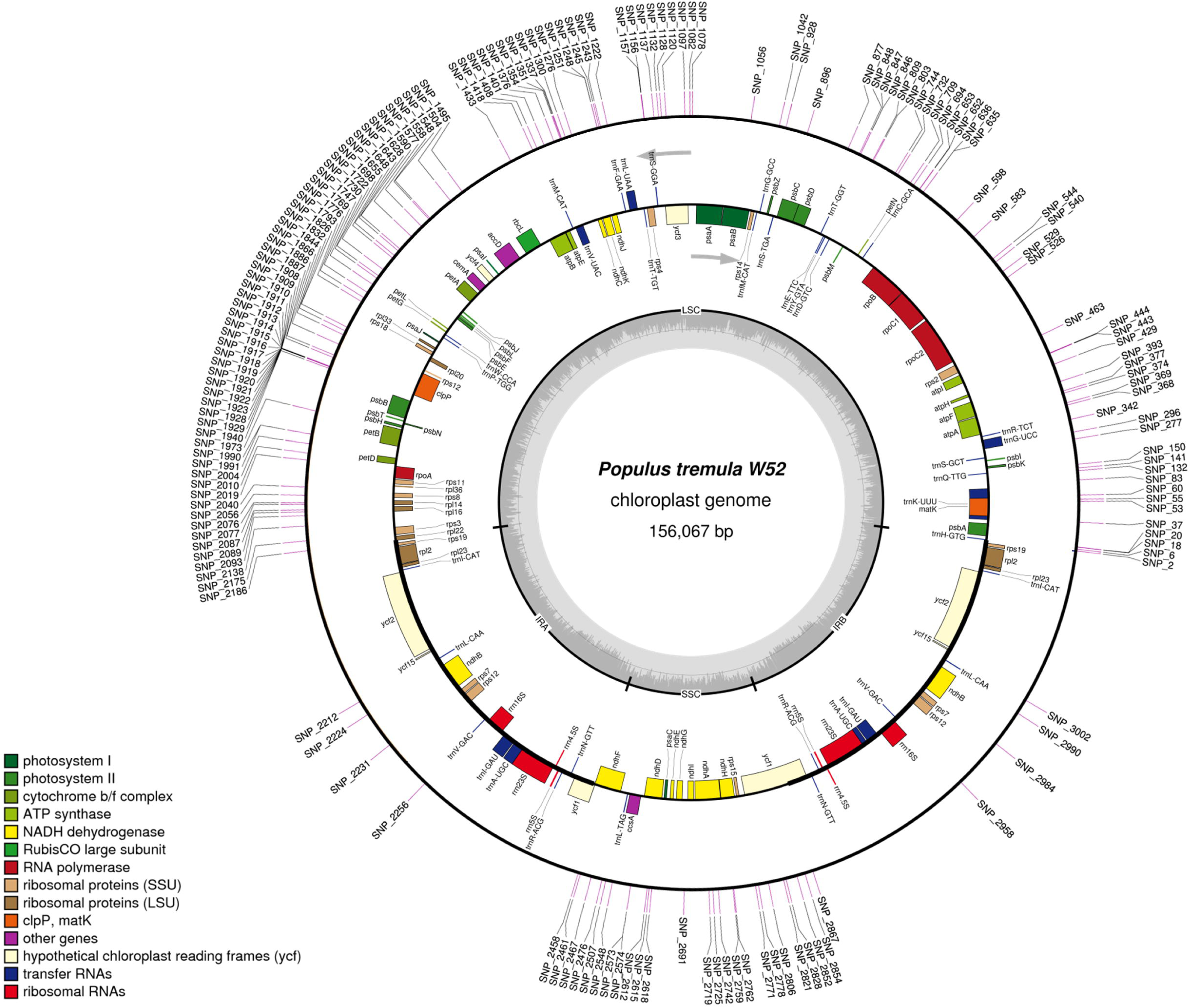
Gene map of the complete *P. tremula* W52 chloroplast genome (GenBank accession number KP861984) and whole-plastome SNP barcode. In total, 163 potential *P*. tremula-specific SNPs (from S9 appendix, “Ptremula_SNPs”; SNPs in InDels were excluded) are depicted on the outer circle as a kind of circular whole-plastome barcode. The middle circle represents the gene map. The grey arrows indicate the direction of transcription of the two DNA strands. A GC content graph is depicted within the inner circle. The circle inside the GC content graph marks the 50% threshold. The mean GC content was found to be 36.76%. The maps were created using OrganellarGenomeDraw [49].

The following genes were predicted in the *P. tremula* chloroplast genome: 85 putative protein-coding genes, 1 pseudogene (*ycf1* in the IRa), 37 tRNA genes as well as 8 rRNA genes. The protein-coding genes *clpP, ycf3*, and *rps12* contained two introns, while 8 additional genes contained one intron each. The *rps12* gene was found to undergo *trans*-splicing, with one exon located in the LSC region and the other two exons located in both IR regions, respectively.

The border sequences between the IR, LSC and SSC regions vary among different species. In *P. tremula*, the borders are located in the *rpl22* gene (LSC-IRa), in the *ycf1* pseudogene (IRa-SSC), in the *ycf1* gene (SSC-IRb) and upstream of the *trnH-GUG* gene (IRb-LSC; Fig. 1). The sequence length difference between the chloroplast genomes of W52 (156,067 bp) and 717–1B4 (156,641 bp) is mainly due to a length difference in the region of the IRa-SSC-border (border at positions 111,876/111,877 of the W52 sequence according to CPGAVAS annotation) that was experimentally confirmed by PCR (S4 appendix) using the primers “Ptre_IRA_SSC_for/rev” (primer sequences in S1 appendix). This length difference is due to a slight expansion of the IRa of 717–1B4 (28,133 bp) compared to W52 (27,509 bp) resulting in an expansion of the truncated *ycf1* gene in the IRa of 717–1B4 compared to W52. The IRa-SSC-border region is also highly variable among the other *Populus* individuals with completely sequenced chloroplast genomes (see next section).

In total, 75 single nucleotide variations, multi nucleotide variations or small InDels were detected in the 717–1B4 chloroplast sequence compared to W52 (S5 appendix; worksheet “variations_cpDNA”) when mapping the 717–1B4 NGS reads to the W52 chloroplast sequence and calling nucleotide variants using CLC GWB. Considering all positions with single or multi nucleotide variations as SNPs, then 59 SNPs were identified corresponding to a mean SNP frequency of 0.378 SNPs/kb in 717–1B4 compared to W52.

### Comparison of the DNA sequences of all completely sequenced *Populus* chloroplast genomes

Complete cpDNA sequences of *P. tremula* W52 (GenBank accession number KP861984) and *P. tremula* x *P. alba* 717–1B4 (KT780870) were aligned together with the complete cpDNA sequences of seven other *Populus* species (see Introduction) and *Salix interior* (NC_024681; [31]), another member of the *Salicaceae* family. Based on this alignment (data not shown), a phylogenetic tree was constructed (Fig. 2). As expected, all *Populus* cpDNA sequences clustered together placing S. *interior* as outgroup. Within the *Populus* cluster, two major clades were resolved and *P. cathayana* appeared as outgroup. Within the upper clade (Fig. 2), members of the section *Populus* (W52, 717–1B4 and *P. alba*) clustered together with *P. yunnanensis*, a member of another section (*Tacamahaca*). The new *P. tremula* cpDNA sequences of W52 and 717–1B4 formed a subclade of latest divergence within the upper clade (Fig. 2). The second main clade comprised two members of the section *Tacamahaca* (*P. trichocarpa* and *P. balsamifera*), not clustering together in a subclade, *P. euphratica* as a member of the section *Turanga* and *P. fremontii* belonging to the section *Aigeiros*.

**Fig. 2.**
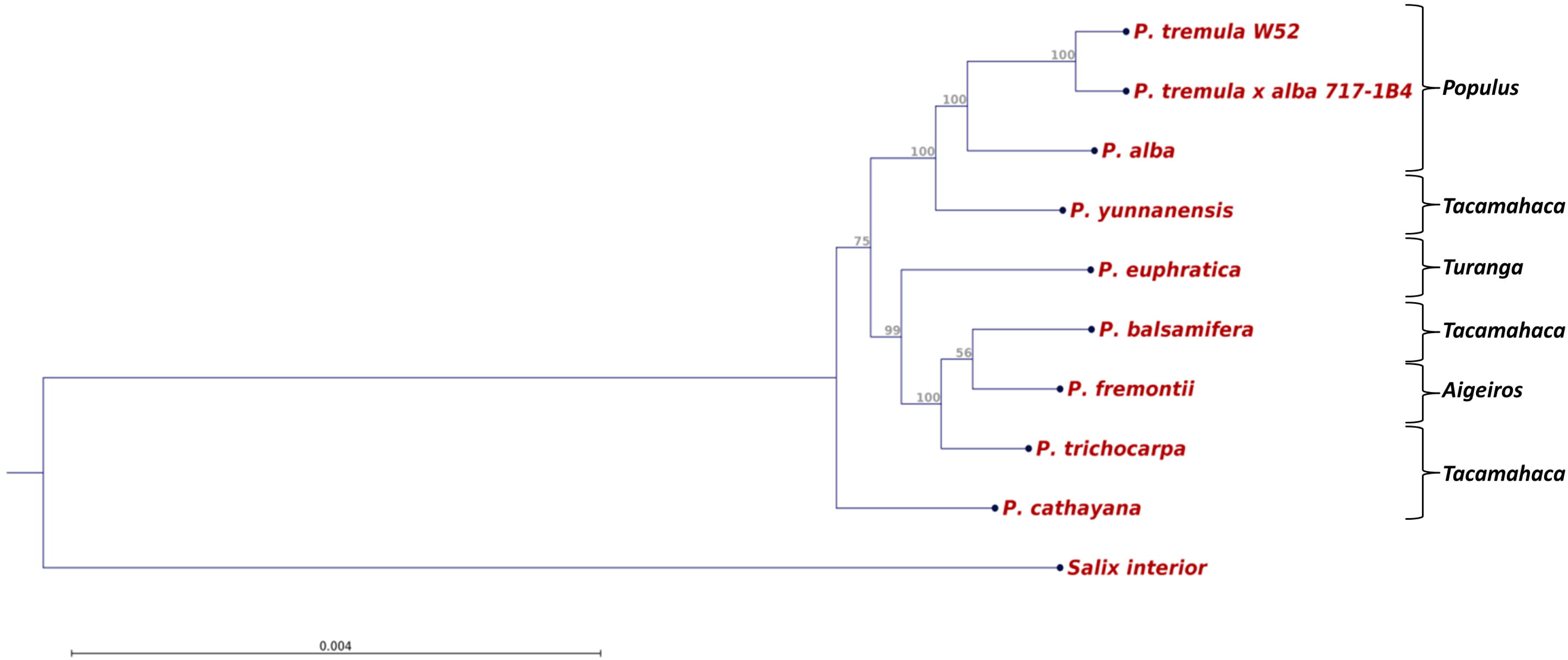
Phylogenetic tree based on all available complete cpDNA sequences of members of the genus *Populus* and of *Salix interior* as a member of another genus of the *Salicaceae* family. The tree was constructed using UPGMA as construction method. Bootstrap support values (%) are shown above branches. GenBank accession numbers: *P. tremula* W52 (KP861984), *P. tremula x P. alba* 717–1B4 (KT780870), *P. trichocarpa* (EF489041), *P. alba* (AP008956), *P. balsamifera* (KJ664927), *P. cathayana* (KP729175), *P. euphratica* (KJ624919), *P. fremontii* (KJ664926), *P. yunnanensis* (KP729176), S. *interior* (NC_024681).

An additional alignment including all complete *Populus* chloroplast sequences (without S. *interior*) was performed to compare the different *Populus* chloroplast genomes with each other (alignment fasta in S6 appendix; consensus sequence in S7 appendix). When comparing the chloroplast genome sequences of W52 with all other *Populus* chloroplast sequences, the 717–1B4 chloroplast sequence is, as expected, the most similar (98.99% nucleotide identity), followed by the *P. alba* chloroplast sequence (98,74%; similarity matrix in S8 appendix).

Based on the alignment of all complete *Populus* chloroplast sequences (S6 appendix), 3,024 positions were called by the SNiPlay tool [47,48] (S9 appendix) which showed DNA sequence variations (SNPs or InDels) in at least one of the chloroplast genomes in comparison to the consensus sequence of all *Populus* chloroplast sequences (in S7 appendix). All variable positions with the related variation statistics as well as the genotyping results of all *Populus* individuals at each variable position are available in S9 appendix (“snp_stats” and “genotyping_data”).

The following regions of the *Populus* chloroplast consensus sequence showed the highest frequencies of variations among all sequenced *Populus* chloroplast genomes (considering intervals of 10,000 bp): 1–10,000 bp with 369 variable positions (top1), 110,000–120,000 bp with 327 positions (top2) and 60,000–70,000 bp with 297 positions (top3). The top1-region is located at the 5-prime part of the LSC (from *trnH* to *atpA;* Fig. 1). The alignment of all *Populus* chloroplast sequences in the most 5-prime part of the top1-region (position 1–480 of the *Populus* consensus sequence) including the *trnH-psbA* linker is presented in S10 appendix. The top2-region includes and surrounds the IRa-SSC border (from the *rrnR* gene cluster to ccsA; Fig. 1). The alignment of all *Populus* cpDNA sequences in the border region is shown in S11 appendix. The top3-region is located in the LSC (from *accD* to *rpl33;* Fig. 1).

Filtering all 3,024 variable positions (S9 appendix; “genotyping_data”) for positions showing alleles that occur only in the two genotypes with *P. tremula* chloroplast genomes (W52, 717–1B4) and not in the chloroplast genomes of the other *Populus* individuals representing seven other species, 232 variable positions remained as potential *P. tremula*-specific cpDNA variations (S9 appendix; “Ptremula_snps”). These positions included 163 SNPs that were displayed as a kind of whole-plastome barcode on the W52 chloroplast sequence (Fig. 1, outer circle). Interestingly, the IRa and IRb included only a few potential *P. tremula*-specific SNPs, whereas large clusters of SNPs were detected in the LSC and SSC (Fig. 1).

The *trnH-psbA* linker mentioned above (S10 appendix; position 78–340 bp of the consensus sequence) includes 3 potential *P. tremula-specific* SNPs at positions 160 (G/A), 312 (A/G) and 324 (T/C) as well as an InDel (6bp) at the border between the *trnH-psbA* linker and the *psbA* gene (position 343). These nucleotide variations were further validated by Sanger sequencing of PCR amplicons of the *trnH-psbA* linker region in an extended set of *Populus* individuals comprising 13 *Populus* species belonging to four *Populus* sections (Table 1). All four nucleotide variations were specific to *P. tremula* when compared to the individuals of the other 12 *Populus* species considered in the analysis (Table 1).

**Table 1.**
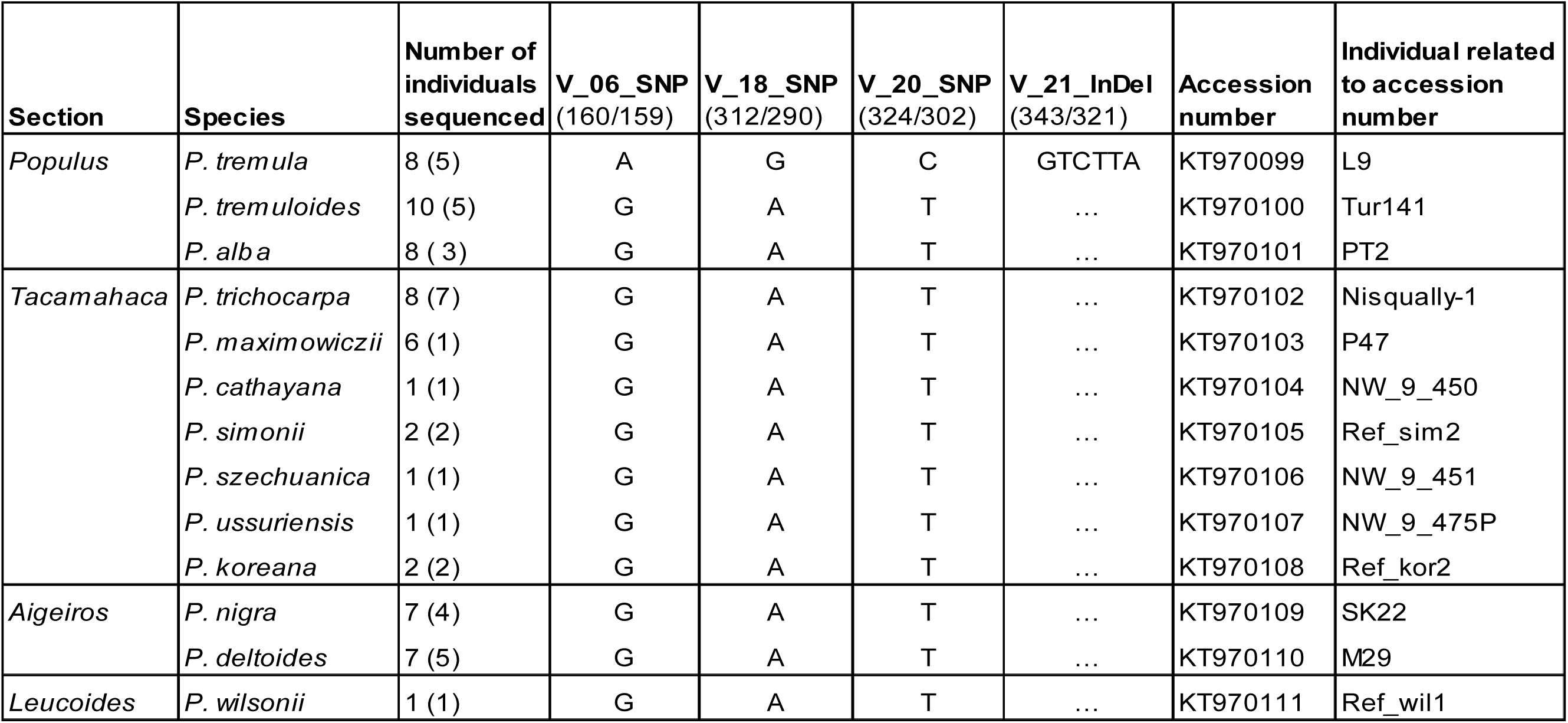
Validation of 4 potential *P. tremula-specific* cpDNA variations in the *trnH-psbA* linker by Sanger sequencing in 13 *Populus* species.

Number of individuals sequenced are given for the 3 SNPs and the InDel (in brackets for the InDel; the InDel region was not readable in all sequences). The numbers of the different variations (V), e.g. V_06, corresponds to the SNP number in S9 appendix (“Ptremula_snps”). The position of the variation in the consensus/*P. trichocarpa* cpDNA sequence is given below the variation number in brackets. GenBank accession numbers are provided for representative Sanger sequences of one individual per species.

### Organization of the *P. tremula* mitochondrial genome and sequence variations between W52 and 717–1B4

The complete DNA sequences of the mitochondrial genomes of the male *P. tremula* clone W52 (GenBank accession number KT337313) and the female *P. tremula* x *P. alba* clone INRA 717–1B4 (KT429213) were assembled using NGS data of total genomic DNA. The mitochondrial genomes are 783,442 bp (W52; Fig. 3) and 783,513 bp (717–1B4) in size and organized as single circles.

**Fig. 3.**
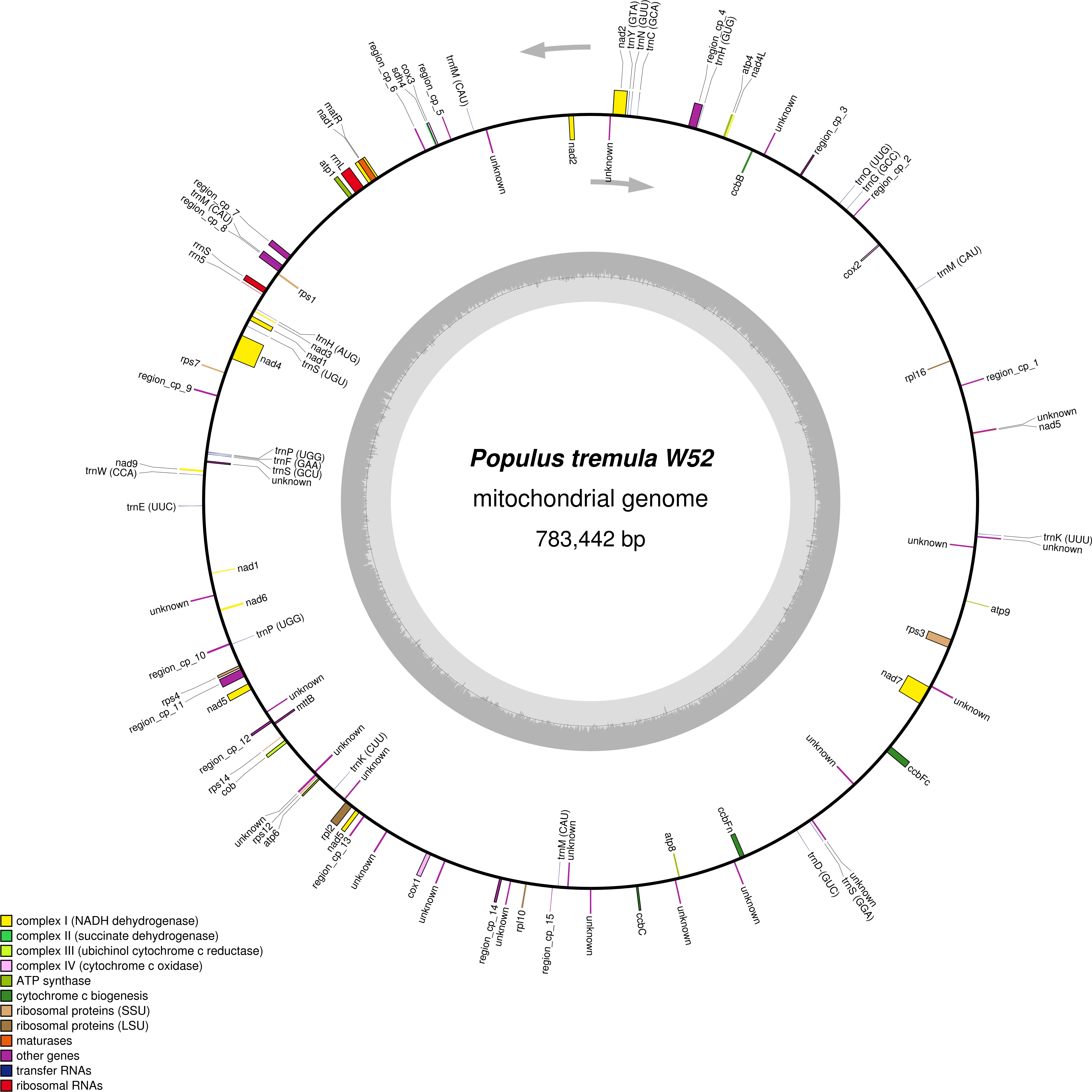
Gene map of the complete *P. tremula* W52 mitochondrial genome (GenBank accession number KT337313). In addition to protein-coding and structural RNA genes, regions of at least 90% similarity to the chloroplast genome of W52 are depicted (“region_cp_1 to region_cp_15”; coloured as “other genes”). The grey arrows indicate the direction of transcription of the two DNA strands. A GC content graph is depicted within the inner circle. The circle inside the GC content graph marks the 50% threshold. The map was created using OrganellarGenomeDraw [49].

Chloroplast-like DNA sequence regions with at least 90% similarity to the W52 chloroplast sequence account for about 2% of the W52 mitochondrial genome and are distributed among 15 distinct regions in the genome (Fig. 3, “region_cp_1” to “region_cp_15”). Most of these regions were also featured by increased coverage (compared to all other regions) when mapping the W52/717–1B4-NGS reads to the related mitochondrial sequences (data not shown). The increased coverage is due to an unspecific mapping of cpDNA-derived reads in addition to mtDNA-derived reads to these regions. Such high-coverage regions and also low coverage-regions (see also Materials and Methods) were validated by Sanger-sequencing of PCR amplicons including (overlapping parts of) these regions (related primer sequences in S1 appendix). The 38 new Sanger sequences were added to an existing dbGSS library of mtDNA Sanger sequences of 717–1B4 at GenBank [12] (dbGSS library accession number LIBGSS_039210; GenBank accession numbers of the new sequences within the library: KS308288 ff.). The mtDNA sequence of 717–1B4 (KT337313) was validated with the Sanger sequences as well as *P. tremula* mitochondrial NGS-scaffolds that were independently generated in the scope of the *P. tremula* v1.0-UPSC genome assembly which is available since recently at PopGenie v3 [76,77]. PopGenie scaffolds were selected by BlastN analysis (using 100,000 bp fragments of the mtDNA sequence as queries) based on similarity to the query (at least 99%) and subject overlap (at least 97%). The Sanger sequences were mapped together with the selected PopGenie scaffolds to the complete mtDNA sequence of 717–1B4 (SeqMan Pro v10.1.2.). The mapping result is presented in S12 appendix and shows that the mtDNA sequence is completely covered by overlapping mapped sequences.

Fig. 3 shows a gene map of the complete *P. tremula* W52 mitochondrial genome. In total, 55 putative protein-coding genes, 22 tRNA genes and 3 rRNA genes were identified. The genes *rps1, rps14, rpl16* and *mttb* are probably pseudogenes. All of the known genes coding for subunits of proteins of the respiratory chain were identified with the exception of *sdh3*, which is beside *sdh4* one of the genes coding for complex II-subunits (Fig. 3). Several genes coding for ribosomal proteins were missing, *i.e., rpl5* (large subunit) as well as *rps2, rps10, rps11, rps13* and *rps19* (small subunit). The genes *nad1, nad2* and *nad5* were predicted to be fragmented in five exons each, belonging to more than one distinct transcription units. The maturation of these genes requires *cis-* as well as trans-splicing events. For the following genes also more than one exon was predicted: *ccbFc, nad4, nad7, rpl2*, and *rps3* (Fig. 3).

A total of 49 nucleotide variations were identified in the mitochondrial genome of 717–1B4 compared to W52 (S5 appendix; “variations_mtDNA”) after mapping the 717–1B4 reads to the W52 mitochondrial sequence and subsequent nucleotide variant-calling by CLC GWB using the same parameters as for the above-mentioned variant detection in the chloroplast genome of 717–1B4. Counting all positions with single or multi nucleotide variations as SNPs resulted in the identification of 51 SNPs. This SNP-number corresponds to a mean SNP frequency of 0.065 SNPs/kb in the mitochondrial genome of 717–1B4 compared to W52.

## Discussion

Poplar breeding has a long history and began in Europe about 80 years ago, presumably with inter-specific hybridization between *P. deltoides*, introduced from northern America, and the native *P. nigra*. Until today, only a few poplar species are being used in these controlled hybridizations. The classical breeding strategy is to establish a first generation (F1) through inter-specific hybridization combined with reciprocal recurrent selection (RRS) of the parental species [28]. Other breeding strategies comprise backcrossing, multiple-species hybridization, polyploidy, and somaclonal variation [28].

Along with the completion of the *P. trichocarpa* reference genome sequence, the first of any tree, in 2006 [23], and further development of the next and third generation sequencing platforms, genomics-based breeding has become a realistic tool in *Populus* breeding. The *Populus* genome includes 41,335 loci containing protein-coding transcripts ([50]; last page view November 9, 2015). However, for most of them the functions are unknown. It is expected that as phenotyping approaches will be more and more coordinated with the sequencing platforms, the functions of most, if not all, genes will be unraveled. For *Populus*, the candidate gene approach is promising as associations to specific marker-traits have been found for many genes [51]. Also, marker-assisted selection (MAS) is feasible today in poplar as a large number of marker-trait associations have been detected [28]. However, candidate gene selection and association genetics are dominated by use of nuclear genes in poplar breeding, and so far neglect the important function of genes located in chloroplast and mitochondrion organelles.

In this study, we present the complete chloroplast and mitochondrial genomic sequence of *P. tremula*, a member of the section *Populus* [20]. The genomes of both organelles were sequenced in the clones *P. tremula* W52 and *P. tremula* x *P. alba* 717–1B4. The male *P. tremula* clone W52 was selected phenotypically on the basis of its growth performance in former East Prussia (Russia). Later, this clone was used as pollen donor in several hybrid aspen breeding programs [37]. The female *P. tremula* x *P. alba* interspecific hybrid (*P. x canescens*) clone 717–1B4 was originally selected in 1992 within a panel of bred *Leuce* poplars (now classified as *Populus*) by Lemoine [38], but not selected for agronomic purpose due to its high susceptibility to *Agrobacterium tumefaciens* infection [52]. Today, it is universally used by scientists worldwide as a model woody species to discover gene expression, gene function and expression localization.

The identified genome sizes of the *P. tremula* chloroplast genomes of W52 (156,067 bp) and 717-1B4 (156,641 bp) are within the size range of all completely sequenced *Populus* chloroplast genomes (see Introduction). The observed variation in size between both genomes was shown to be due to a slight expansion of the IRa (at the IRa-SSC border) in 717–1B4 compared to W52 (S4 and S11 appendices). IRs are one of the hotspots for structural rearrangements within plastid genomes and are frequently subjected to expansion, contraction or even complete loss [18]. It was previously shown that expansions and contractions of IR regions caused variation of size among chloroplast genomes [53].

The identified numbers of potential protein-coding genes (85), pseudogenes (1), tRNA genes (37) and rRNA genes (8; Fig. 1) in the *P. tremula* chloroplast genome are within the distribution range of related numbers in land plants (see Introduction) and similar to other *Populus* chloroplast genomes [17]. All protein-coding genes belonging to the proposed gene content of Angiosperms (putatively ancestral gene content without genes lost during the evolution of Angiosperms) [18] were predicted in the *P. tremula* chloroplast genome, with the exception of *psaM, rpl32*, and *rps16*. These genes are also missing in all other *Populus* chloroplast genomes (mentioned in the Introduction). The genes *rpl32* and *rps16* are also missing in chloroplast genomes of other angiosperm species [54] and *psaM* was shown to be missing in other dicotyledonous plant species [55]. Beyond the proposed gene content of Angiosperms [18], the following genes were predicted in the *P. tremula* chloroplast genome: *psaC, psbB*, and *psbN*. These genes were also predicted in the other *Populus* chloroplast genomes. The gene *psaC* is one of the photosystem I genes that belong to genes shared by both, cyanobacteria and plant chloroplasts [55]. The photosystem II-genes *psbB* and *psbN* are parts of the pentacistronic *psbB* transcription unit that has been well characterized in vascular plants [56].

Since the substitution rate in the DNA of chloroplast genomes is much lower than of nuclear genomes in plants and because of the feature of uniparental inheritance, chloroplast genomes are valuable sources for phylogenetic analyses of higher plants [57]. A phylogenetic tree constructed from all complete *Populus* chloroplast sequences (Fig. 2) was not congruent with the assignment of the species to different *Populus* sections. Although the three members of the section *Populus* clustered together in a subclade, the related main clade included also a member of the section *Tacamahaca* (*P. yunnanensis*). In a phylogenetic analysis of seven *Populus* species from China belonging to the section *Tacamahaca, P. yunnanensis* was shown to be distinct from the other species based on the analysis of nuclear inter-simple sequence repeats [58]. In a recent study on whole-plastome phylogeny of all common poplar species in North America, a deep plastid divergence between two closely related members of the section *Tacamahaca, P. trichocarpa* and *P. balsamifera* was demonstrated [31]. This divergence was also obvious in our study (Fig. 2). Huang *et al*. showed that *P. trichocarpa* and *P. balsamifera* are as different to each other as they are to *P. fremontii* belonging to another section (*Aigeiros*) [31] – a finding that is in line with our results. Another recent study is not directly comparable with our study because it was based on nuclear sequences and plastid fragments [59]. In general, potential sequencing errors in NGS-sequenced complete chloroplast genomes cannot be excluded and may influence the resulting phylogenies. According to Genbank ([60]; last page view November 9, 2015), the chloroplast sequences of *P. cathayana* and *P. yunnanensis* are so far classified as “unverified”.

SNPs and other nucleotide variations in chloroplast genomes that allow for differentiation between *Populus* sections, species groups or species or even identification of the geographic origin of *Populus* trees of a given species are important for practical breeding and research, and are thus highly desirable. Some species-specific cpDNA-markers based on SNPs have been already developed for *Populus* using single-loci for barcode development [11,14], but, so far, no studies on the development of whole-plastome barcodes – according to the recently proposed concept of super-barcoding [61] – are available in *Populus*. The 3,024 variable positions identified in the *Populus* chloroplast consensus sequence when comparing all complete *Populus* chloroplast genomes in this study are a first step towards this direction. These positions can be exploited for the future development of novel markers for species (section) differentiation in *Populus*.

The 5-prime part of the LSC (1–10,000 bp) was identified as the chloroplast region with the highest frequency of variable positions (top1-region) when considering the distribution of all 3,024 variable positions in the *Populus* cpDNA consensus sequence. The intergenic linker *trnH-psbA, psbK-psbI* and *trnG-psbK* included in the top1-region showed the highest percentage of variable sites when comparing 20 chloroplast regions in seven *Populus* species [13]. Also in a recent comparison of 20 chloroplast regions in 14 poplar species, some intergenic linkers of the top1-region were among the linkers with highest numbers of species-specific SNPs and InDels (*matK-trnK, trnH-psbA, psbK-psbI;* [14]).

In total, 163 potential *P*. tremula-specific SNPs were identified in the chloroplast genomes of W52 and 717–1B4. These SNPs were presented as a whole-plastome barcode on the *P. tremula* W52 cpDNA sequence (Fig. 1). This barcode includes several SNP clusters in the LSC and SSC but only a few SNPs in the IRs. This finding is in line with previous studies [62,63]. The SNP rate in IR regions was 4 times lower than that in the single copy regions in a comparative study on cpDNA sequences of two rice varieties [62]. In another study, the synonymous substitution rate of IR regions was roughly 5 times lower than that of the single copy regions when the chloroplast genome sequences were compared among different species [63]. Out of the 163 potential *P. tremula*-specific SNPs, three SNPs together with one InDel in the *trnH-psbA* linker were validated by Sanger sequencing in an extended set of *Populus* individuals of different species (Table 1). These four nucleotide variations allow for differentiating *P. tremula* individuals from individuals of other members of the section *Populus* (*P. tremuloides, P. alba*)*, Tacamahaca* (*P. triochocarpa, P. maximowiczii*) and *Aigeiros* (*P. nigra, P. deltoides*) and probably also of *Leucoides*.

Mitochondria are very important for plant breeders because of a trait called cytoplasmic male sterility (*cms*). Plants displaying *cms* are male sterile, *i.e*., they do not form viable pollen, thus allowing for establishing hybrid seed stocks. The *cms* trait is correlated with distinct rearrangements in the mtDNA and, thus, maternally inherited with mitochondria. The recent sequencing of the entire mitochondrial genome of *Hevea brasiliensis, e.g*., revealed a rearrangement in the genome resulting in a novel transcript that contained a part of *atp9* and caused male sterility through a slight reduction of ATP production efficiency in the analyzed male sterile tree [64]. An interrelationship to the nucleus exists, *i.e*., nuclear genes can overcome the mtDNA rearrangements and restore fertility in the *cms* plants. Another important trait associated with plant mitochondria is oxidative burst involved in apoptotic responses including programmed cell death [65]. Apoptosis is induced by reactive oxygen species (ROS), and interestingly, mitochondria are both source and target of ROS [66]. Programmed cell death in plants is correlated with multiple defense functions, *e.g*., the hypersensitive response (HR) to pathogens [67].

With the complete nucleotide sequences of the mitochondrial genome of the clones *P. tremula* W52 and *P. tremula* x *P. alba* 717–1B4, we provide the first in the genus *Populus* and even in the family of *Salicaceae*. In the order of *Malpighiales*, only two mitochondrial genomes were published before: *Ricinus communis* (GenBank accession number HQ874649; [68]) and *Hevea brasiliensis* (AP014526; [64]). The genome sizes of the provided *P. tremula* genomes (783,442 bp in W52 and 783,513 bp in 717–1B4) are very similar and range between the genome sizes of the mitochondrial genomes of *R. communis* (502,773 bp) and *H. brasiliensis* (1,325,823 bp). Tuskan *et al*. reported on a putative master molecule of 803 kb for the mtDNA of *P. trichocarpa* [23] which is a bit larger than the *P. tremula* mtDNA that we assembled (783 kb). No statement on a finishing of the master molecule (e.g., by circularization) is given by the authors. A finishing may result in a shorter sequence. Moreover a general size difference between the mtDNA sequences of *P. tremula* and *P. trichocarpa* that represent species of two different *Populus* sections, cannot be excluded.

Critical connections of the *P. tremula* mtDNA sequences were validated by Sanger sequencing (GenBank dbGSS library accession number LIBGSS_039210) and the 717–1B4 mtDNA sequence could be fully covered in a validation mapping of the Sanger sequences together with mitochondrial *P. tremula* NGS-scaffolds from PopGenie [76,77] (S12 appendix). Nevertheless, single sequencing errors caused by NGS cannot be fully excluded.

In general, the majority of the mitochondrial genes code for important proteins of the electron transport chain including oxidative phosphorylation processes, *e.g*., cytochrome c oxidase, ATP synthase and NADH dehydrogenase. Other genes code for structural tRNA and rRNA molecules. In total, 80 genes were identified in the *P. tremula* mitochondrial genome, including 55 protein-coding genes, 22 tRNA genes and 3 rRNA genes (Fig. 3 for W52). This total gene number is in the distribution range of related numbers in land plants (see Introduction).

Mitochondria are characterized by high genome reorganization, rearrangement and/or horizontal gene transfer [69,70], *e.g*., chloroplast and nuclear sequences have been found in mitochondrial genomes or *vice-versa*, mitochondrial sequences in the nuclear genome. These are ongoing processes in plants. Most of the transfers in angiosperms involve ribosomal protein genes [71]. Thus, it is not unexpected that 6 of the ribosomal genes belonging to the ancestral gene content of the mitochondrial genome of flowering plants [19] were not predicted in *P. tremula*. The genes *rps2* and *rps11* missing in the mitochondrial genome of *P. tremula* are also lacking in the mitochondrial genomes of *R. communis* and *H. brasiliensis* [64,68]. Other ribosomal genes missing in *P. tremula* (*rps10* and *rps19*) are also not present in *H. brasiliensis*, but are present in *R. communis*. The ribosomal genes *rpl5* and *rps13* missing in *P. tremula* are present in the mitochondrial genomes of *H. brasiliensis* and *R. communis* [64,68]. The *rpl5* gene is lacking from many of the sequenced plant mitochondrial genomes [45]. The absence of the *rps13* gene from mitochondrial genomes has been shown for many members of the rosids subclass [72]. Apart from ribosomal proteins, the loss of two respiratory genes, *sdh3* and *sdh4* (encoding subunits 3 and 4 of succinate dehydrogenase) has been reported from the mitochondrial genome of various angiosperms [71]. Whereas *sdh4* could be annotated in the *P. tremula* mitochondrial genomes, *sdh3* is lacking in *P. tremula* in contrast to *R. communis* and *H. brasiliensis*, where both genes could be annotated in the mitochondrial genomes [64,68].

It is generally accepted that for land plants the plastid and nuclear genomes have an approximately three-to tenfold greater mutation rate than the mitochondrial genome [19,73], with some exceptions [74]. We identified 59 SNPs (0.378 SNPs/kb) in the chloroplast genome and 51 SNPs (0.065 SNPs/kb) in the mitochondrial genome of 717–1B4 compared to the related W52 reference genomes. Thus, the mean SNP frequency was found to be nearly six fold higher in the chloroplast genome than in the mitochondrial genome when considering both individuals sequenced in this study.

Due to the comparatively low mutation rate in mitochondrial genomes, barcoding attempts based on mtDNA are rare in land plants. The first mitochondrial markers for species-differentiation in *Populus* based on SNPs were developed recently [12] and the described *P. tremula-specific* SNPs were also obvious in the *P. tremula* mitochondrial genomes sequenced in this study. An attempt to also differentiate between cultivars by mitochondrial SNPs has been made recently in date palm [75]. The authors identified 188 SNPs among nine cultivars (18–25 SNPs in each cultivar compared to the reference sequence) in the mitochondrial genome. Most SNPs were shared among cultivars with only 14 unique to a single cultivar [75]. Thus, the authors recommended using nuclear SNPs instead of mitochondrial SNPs for molecular characterization of date palm cultivars. Whether mitochondrial barcodes are suitable for the differentiation of poplar cultivars remains to be tested in future studies.

With the availability of whole genome cpDNA and mtDNA sequences of *P. tremula*, marker-assisted poplar breeding and association genetics can profit from the knowledge of these DNA-containing cellular compartments. Besides the complete nuclear genome sequence of *P. trichocarpa* [23], other nuclear genome sequences from additional *Populus* species will become available in the forthcoming years, *e.g., P. tremula* and *P. tremuloides* [76,77]. Combined with classical poplar breeding approaches, genomic breeding considering the three DNA-containing cellular compartments is a first step towards a holistic approach in *Populus* breeding.

## Acknowledgements

We acknowledge the support by CEA-IG/CNG performing the DNA QC in its DNA and Cell Bank service and providing access to their Illumina Sequencing Platform. We are grateful to the teams of Anne Boland (DNA and Cell Bank service) and MarieThérèse Bihoreau (Illumina Sequencing and Infinium genotyping facilities). We thank Isabelle Le Clainche and Aurélie Chauveau for performing DNA extraction and sequencing. We are grateful to Katrin Groppe, Manuela Will, and Susanne Bein for technical assistance and to Dina Führmann for language editing.

## Supporting Information

**S1 appendix. Primer sequences for PCR amplification of specific DNA regions in the cpDNA and mtDNA of *Populus***. DNA sequences of cpDNA primers *trnH-psbA* and *rpoC2-rpoC1* were taken from [13]. DNA sequences of mtDNA primers were derived from the mtDNA sequence of 717–1B4.The location of the mtDNA primers, the primer combinations used for the generated amplicons and the theoretically/experimentally length of the amplicons are provided in addition. In case of primers with high similarity (100% identity or 1 mismatch) to the chloroplast genome, these primers were combined only with primers that did match to the mtDNA but not to the cpDNA.

**S2 appendix. Testing of *P. tremula* x *P. alba* 717–1B4 (sample P1) with the *Populus* chloroplast PCR-RFLP markers *rpoC2-rpoC1* (*Mls*I) and *trnH-psbA* (*Alw*26I) [13,14]**. Three other *P. tremula* individuals (W52, Graupa 103, GD5) and three *P. alba* individuals (6K6, Villa Franka, 14P11) were used as controls.

**S3 appendix. Protocol for trimming and assembly of the NGS data**.

**S4 appendix. PCR amplicons of the IRa-SSC-border region in *P. tremula* W52, *P. tremula* Br11 and *P. tremula* x *P. alba* 717–1B4**. DNA sequences of the primers “Ptre_IRA_SSC_for/rev” are given in S1 appendix. The theoretical lengths of the PCR products are 665 bp (W52) or 1126 bp (717–1B4), respectively, based on the related cpDNA sequences.

**S5 appendix. Nucleotide variations in cpDNA and mtDNA of 717–1B4 compared to W52**. Nucleotide variations (SNPs and small InDels) in cpDNA (worksheet “variations_cpDNA”) and mtDNA (“variations_mtDNA”) of 717–1B4 compared to W52 (reference). The variations were identified using CLC GWB.

**S6 appendix. Alignment fasta for the alignment of all available complete *Populus* cpDNA sequences**.

**S7 appendix. Consensus sequence (fasta) based on the alignment of all available complete *Populus* cpDNA sequences**.

**S8 appendix. Similarity matrix of all *Populus* available complete cpDNA sequences. Identities (in %) of the cpDNA sequences in pairwise comparisons**.

**S9 appendix. Nucleotide variations between all available complete *Populus* cpDNA sequences**. The variations were identified by SNiPlay (pipeline v2; [47]). Positions are related to the *Populus* chloroplast consensus sequence (S7 appendix). Nucleotide variation statistics in all *Populus* cp sequences (worksheet “snp_stats”) and related genotyping of all analyzed *Populus* individuals at each variable position of the *Populus* chloroplast consensus sequence (“genotyping_data”). Selection of variations that occur in the *P. tremula* cpDNAs of W52 and 717–1B4 but not in the cpDNAs of all other sequenced *Populus* chloroplast genomes (“Ptremula_snps”).

**S10 appendix. Alignment of all available complete *Populus* cpDNA sequences in a section of the top1-region**. This section represents the broader *trnH-psbA* linker region.

**S11 appendix. Alignment of all available complete *Populus* cpDNA sequences in a section of the top2-region**. This section represents the broader IRa-SSC linker region.

**S12 appendix. Alignment of all Sanger sequences of 717–1B4 mtDNA and mitochondrial *P. tremula* scaffolds from PopGenie [76,77] to the mtDNA sequence of 717–1B4 (GenBank accession number KT429213)**. The alignment was created by SeqMan Pro (v10.1.2. DNA Star, Madison, USA). The figure gives the alignment overview (“strategy view” of SeqMan Pro). Sanger sequences of PCR amplicons of 717–1B4 are named with the numbers of the forward/reverse primer, e.g. “717–1B4_1834”_for. *P. tremula* or *P. tremula* x *P. tremuloides* (T89) scaffolds selected from PopGenie are named with “Potra…” or “Ptrx…”, respectively. Forward and reverse Sanger sequences that did not overlap were combined and separated by respective N-stretches.

